# Single Cell Observations Show Persister Cells Wake Based on Ribosome Content

**DOI:** 10.1101/247221

**Authors:** Jun-Seob Kim, Thomas K. Wood

## Abstract

Since persister cells survive antibiotic treatments through dormancy and resuscitate to reconstitute infections, it is imperative to determine the rate at which these cells revive. Using two sets of *Escherichia coli* persister cells, those arising naturally at low levels and those generated at high levels by ceasing transcription via rifampicin pretreatment (shown to be bona fide persisters through seven sets of experiments), we used microscopy of single cells to determine that persisters have low levels of antibiotic-corrupting proteins and that their resuscitation is heterogeneous and includes cells that grow immediately. In all, five phenotypes were found for persister cell resuscitation: (i) immediate division, (ii) immediate elongation followed by division, (iii) immediate elongation but no division, (iv) delayed elongation/division, and (v) no growth. In addition, once cell division begins, the growth rate is that of exponential cells. Critically, the greater the ribosome content, the faster the persister cells resuscitate.

## INTRODUCTION

Bacteria have multiple defense mechanisms against environmental stresses (Requena 2012) and one of most important is bacterial persistence. Similar to resistance, persister cells survive antibiotic exposure (Kim *et al.*, 2011, Lewis 2010). However, unlike resistant cells, persisters do not inherit the antibiotic tolerance and do not grow in presence of antibiotics. Hence, upon adding fresh medium, persisters grow like the parental population (Keren *et al.*, 2004, Wiuff *et al.*, 2005). Persister cells are non-growing cells (Bigger 1944, Hobby *et al.*, 1942, Kwan *et al.*, 2013, Shah *et al.*, 2006) that form from a depletion of protein production capacity; i.e., a cessation of transcription, translation, and ATP production (Cheng *et al.*, 2014, Conlon *et al.*, 2016, Dorr *et al.*, 2010, Kwan *et al.*, 2013); however, some groups also have reported that they are not completely dormant (Orman and Brynildsen 2013). Hence, the cellular processes of persisters are not corrupted by antibiotics due to their reduced metabolism (Lewis 2007). Persisters most likely form as a stress response such as that induced by antibiotics (Dorr *et al.*, 2010, Kwan *et al.*, 2013, Van den Bergh *et al.*, 2016); however, they also have been reported to arise spontaneously (Balaban *et al.*, 2004).

Most microorganisms experience stress since their environmental conditions change (e.g., nutrient levels); hence, dormancy or reduced metabolism is a very common response to environmental stress. One concept in microbial ecology is that these cells with reduced metabolism form a seed bank to give cells a chance to revive after stress and ultimately, provide diversity in microbial communities (Lennon and Jones 2011) and a stable pool of microbial communities (Stevenson 1977). This concept is supported by the many groups that have observed inactivated cells in various environments (Lennon and Jones 2011). Furthermore, persister cells appear to be part of the low-metabolism continuum that includes viable but non-culturable (VBNC) cells (Ayrapetyan *et al.*, 2015), those cells that form from starvation. In our hands, the non-dead cells that form under VBNC-inducing conditions have the same phenotypes as persister cells (Kim *et al.*, 2017). In medicine, persister cells are thought to be responsible for some recurring infections (Fisher *et al.*, 2017). Hence, persister cells should be prevalent both in medicine and in the environment (since all nearly all cells experience starvation conditions).

Many studies have been performed investigating the mechanism of persister cell formation (Dorr *et al.*, 2010, Harrison *et al.*, 2009, Kim and Wood 2010, Korch *et al.*, 2003, Kwan *et al.*, 2013) and regarding eradication of persister cells (Chowdhury *et al.*, 2016, Kwan *et al.*, 2015); however, there are only few publications on persister resuscitation (Balaban *et al.*, 2004, Joers *et al.*, 2010). Using a microfluidic device, Balaban and coworkers found *Escherichia coli hipA7* persister cells wake in approximately one hour and used stationary-phase cells to estimate waking occurs in 16 h (Balaban *et al.*, 2004). The Tenson group also used stationary-phase cells of *E. coli* and estimated persister cells start dividing in one to two hours after contact with fresh medium (Joers *et al.*, 2010). Unfortunately, stationary phase cells consist of antibiotic-sensitive cells, tolerant cells, and persister cells, and persister cells are the smallest population. Furthermore, persister cells are not identical to stationary-phase tolerant cells based on transmission electron microscopy which shows a cell structure for each cell type (Cho *et al.*, 2017). These results also illustrate the need for persister cell populations in which persister cells are the dominant cell type in order to more accurately estimate the persister cell wake time.

Here, we observed persister cells on solid cultures and found morphological heterogeneity upon persister cell resuscitation. We used rifampicin-pretreatment to create persister cells (Kwan *et al.*, 2013) since rifampicin pretreatment creates a large number of persister cells (up to 80% of total cells) and examined single cells as they revived. To corroborate the phenotype of the rifampicin-induced persisters, we observed the waking of natural persister cells (those few cells remaining after ampicillin treatment) and confirmed that the same morphological phenotypes were observed upon waking; hence, rifampicin-induced persisters are equivalent to natural persister cells. Using labeled penicillin, we found cells become persisters when they do not contain residual antibiotics; in contrast, dead cells contain high levels of antibiotic. Furthermore, we found that that persister cell resuscitation can occur immediately for persister cells when they contact a carbon source and that this re-growth consists of either cell elongation followed by division or division without elongation; some cells also had a delay in elongation and division. In addition, by using an *E. coli* strain that correlates ribosome content via the green fluorescent protein (GFP), we found that the heterogeneity upon persister waking is caused by differences in ribosome content prior to antibiotic treatment: cells with more ribosomes wake faster.

## MATERIALS AND METHODS

### Bacterial strain and growth conditions

*E. coli* K-12 BW25113 (Baba *et al.*, 2006) and *E. coli* K-12 MG1655-ASV (*rrnb*P1::GFP[ASV]) (Shah *et al.*, 2006) were used in this study. All experiments were conducted at 37 ºC in Luria-Bertani (LB) medium (Sambrook *et al.*, 1989), and planktonic cultures were shaken at 250 rpm in 250 mL Erlenmeyer flasks.

### Persister cell viability assay (planktonic cells)

Overnight cultures of BW25113 were inoculated into 25 mL fresh LB using a 1:1000 dilution and incubated until the turbidity at 600 nm reached to 0.8. Rifampicin (100 µg/mL) was added for 30 min to induce persister cell formation (Kwan *et al.*, 2013). The Rif-treated culture was harvested by centrifugation at 3,500 × *g*, 4 ºC for 10 min (note the cold centrifugation step did not affect persistence, data not shown). The cell pellet was suspended into 25 mL pre-warmed fresh medium containing ampicillin (100 µg/mL) and incubated for 3 hour to remove the non-persister cells (Kwan *et al.*, 2013). Every hour during ampicillin treatment, 1 mL of cell culture was washed and diluted serially by PBS. Samples (10 µL) were spotted onto LB agar plates, and colonies were counted after overnight incubation at 37 ºC to determine the number persister cells.

For the assay for natural, planktonic persisters, ampicillin (100 µg/mL) was added to exponentially-grown bacterial cultures (turbidity at 600 nm ~ 0.8). Every hour during ampicillin treatment, 1 mL of cell cultures was washed and diluted serially by PBS. Samples (10 µL) were spotted onto LB agar plates, and colonies were counted after overnight incubation at 37 ºC to determine the number persister cells.

### Persister resuscitation on agarose gel pads

Low melting agarose (1.5 %) (Nusieve GTG agarose – BMB # 50081) was added to LB medium and melted by microwaving (30 sec at 1,500 Watts). The gel pad template **(Supplementary Figure S1)** was made by placing three microscope glass slides (75 × 25 × 1 mm) side-by-side to form the base with two slides on top of these to form a groove, then the melted LB agarose was poured until it filled the groove. The pad was covered by with another slide glass and held together with 50 g of weight for 30 min to solidify. The slide glass was removed after solidification, and the gel pad was cut to the appropriate size with a gel knife and moved to a new slide glass. After persister cells were added, the, agarose gel was covered by a glass coverslip and incubated for 20 min at room temperature to dry. The edge of the coverslip was sealed by nail polish to prevent evaporation and drying of the agarose pad during long observations. The agarose gel pad with cells were observed by light microscope (Zeiss Axioscope.A1, bl_ph channel at 1,000 ms exposure) every 20 min up to 24h. The microscope was placed in a vinyl glove box (Coy Labs) and warmed by an anaerobic chamber heater (Coy Labs, 8535-025) to maintain 37 ºC.

For the rifampicin-induced persisters, 1 mL of cell cultures were harvested by centrifugation (3,500 × g, 4 ºC for 10 min) and re-suspended into 1 mL of PBS. To remove dead cells and concentrate the natural persisters, cultures were harvested by centrifugation at low speeds (200 × *g*, for 10 min), and the supernatant (containing dead cell and cell debris) was decanted gently. The cell pellet was resuspended in one mL of PBS and harvested again by centrifugation (50 × *g*, for 10 min). The suspension was decanted gently using a pipette (900 µL), the remaining liquid (100 µL, containing persister cells) was concentrated by centrifugation (17,000 × *g* for 10 min), and the pellet resuspended in 5 µL of PBS.

For the GFP-tagged, rifampicin-induced persisters of *E. coli* K-12 MG1655-ASV, the 20 min-drying step was removed. GFP intensity was monitored using a fluorescence microscope (Zeiss Axioscope.A1, bl_ph channel at 1,000 ms exposure and GFP channel at 10,000 ms exposure). As a control, rifampicin-induced persister cells of *E. coli* K-12 MG1655-ASV (30 min pretreatment with 100 µg/mL) were plated onto LB agarose gel pads containing ampicillin (100 µg/mL), and the GFP signal changes were monitored.

### Metabolic activity of persister cells

For exponential cells and rifampicin-induced persister cells, the metabolic activity was measured by flow cytometry. Cells were pelleted and washed twice with PBS. For the dead cells (negative control), 1 mL of exponential culture was centrifuged, resuspended in 70% isopropanol, and incubated for 1 h at room temperature. Propidium iodide (PI, Molecular Probes, Eugene, OR, USA) and redox sensor (BacLight™ RedoxSensor™ Green Vitality Kit, Thermo Fisher Scientific Inc., Waltham, MA, USA) were used with samples incubated at 37 ºC for 10 min with light protection.

The fluorescence signal was analyzed by flow cytometry (Beckman Coulter FC500) using the FL1 and FL3 channels.

### MIC test

Overnight cultures of BW25113, rifampicin-induced persister cells (without ampicillin treatment) and re-grown populations from rifampicin-induced persisters were diluted into fresh LB at 10^4^ cells/mL. Cells (196 µL) were placed in 96-well plates, and ampicillin and Bocillin-FL were diluted and added in 4 µL. The 96-well plates were incubated at 37 ºC overnight, and the concentration at which cells did not grow was used as the MIC.

### Resistance test for rifampicin-induced persister cells

Exponential cells (for natural persisters) and rifampicin-induced persisters were treated with ampicillin (100 µg/mL) for 3 h to remove the non-persister cells. Cells (1 mL) were washed with PBS and re-inoculated into 25 mL of fresh LB and incubated at 37 ºC with shaking (250 rpm) overnight. The re-grown populations were inoculated into 25 mL of LB and incubated until the turbidity at 600 nm reached 0.8. Ampicillin (100 µg/mL) was added to each re-grown population for 3 hour, and the cells were observed for lysis.

### Ribosomal RNA synthesis via flow cytometry analysis and cell sorting

The rifampicin-induced persister cells (*E. coli* K-12 MG1655-ASV) were sorted based on GFP intensity. GFP intensity was analyzed with a Beckman-Coulter Moflo Astrios EQ instrument (Beckman Coulter Life Sciences, Brea, CA, USA). Before the FACS analysis, PI was added to the cell suspension (1.83 mM) to allow the dead cells to be removed based on PI signal (red fluorescence) of persisters compared to live cell (exponential cell) and dead cell control (treated 70% isopropanol for 1 hour to exponential cell). The cell population was determined by forward and side scatter (FSC and SSC), and GFP fluorescence was analyzed with a blue laser (488 nm) and a 513/26 nm bandpass filter (GFP) while PI fluorescent was analyzed with a 620/29 nm bandpass filter. To reduce the time in the PBS buffer, sorted cells were deposited directly onto the LB agarose gel pad on the slide glass (5,000 events in 2 min) before covering with a glass cover slip (to maintain the high concentration of cells in the sorted spot). Sorted cells were observed immediately using a fluorescence microscope (Zeiss Axioscope.A1, bl ph channel at 1,000 ms exposure and GFP channel at 10,000 ms exposure).

### 70S ribosome (active ribosome) purification

70S ribosomes were isolated as described previously with slight modifications (Rivera *et al.*, 2015). Briefly, exponential, rifampicin-induced persister, and stationary phase cultures were chilled immediately on ice-ethanol for 5 min, then harvested by centrifugation at 3,500 × g for 10 min; the cell pellets were resuspended in 13 mL of ribosome buffer (20 mM Tris, 50 mM MgOAc, 100 mM NH4Cl, 0.5 mM EDTA, pH 7.5) and 6 mM β-mercaptoethanol. Cells were disrupted by a French press at 16,000 psi twice, then treated with 5 U of DNase I (RNase-free) on ice for 20 min. Crude cell debris was removed by centrifugation at 30,000 × g for 30 min. To an ultracentrifuge tube with 13 mL of ribosome buffer containing 32% sucrose, 13 mL of cell supernatants were layered carefully. The 70S ribosomes were pelleted by ultracentrifugation at 100,000 × g for 16 hours at 4 ºC. The ribosome pellets were re-suspended in PBS buffer, and the 70S ribosomes were quantified by absorbance at 260 nm (for rRNA).

### Penicillin binding proteins via Bocillin-FL

Exponential cells (turbidity at 600 nm ~ 0.8), stationary-phase cells (turbidity ~ 5), and rifampicin-induced persister cells were incubated with Bocillin-FL (Thermo Fisher Scientific Inc, Waltham, MA, USA) for 3 hours and incubated at 37 ºC in the dark by adding Bocillin-FL to cells in the growth medium for the exponential and stationary cells and by adding it along with ampicillin (100 µg/mL) for the rifampicin-induced persister cells. The samples were harvested by centrifugation at 17,000 × g for 1 min and washed by PBS twice. Samples (5 µL) were observed via fluorescence microscopy (Zeiss Axioscope.A1, bl_ph channel at 1,000 ms and BODIFY channel at 10,000 ms exposure).

## RESULTS

### Rifampicin-induced persisters are bona fide persister cells

To investigate persister cell resuscitation in fresh medium, the rifampicin-induced persisters were employed initially since they are formed in high numbers (Kwan *et al.*, 2013) which enables us to study easily at the single-cell level. As expected, the persister frequency after rifampicin treatment showed a 10^5^ fold increase compared to persister frequency of exponential cells without rifampicin treatment (**Supplementary Figure S2**). The persister cells induced by rifampicin were isolated from non-persister cells by ampicillin treatment to remove the non-persister cells since the non-persisters cells lyse (Kwan *et al.*, 2013).

Along with the tolerance to ampicillin (**Supplementary Figure S2**), we originally demonstrated these rifampicin-induced persister cells are tolerant to ciprofloxacin (Kwan *et al.*, 2013); hence, like natural persister cells (Lewis 2010), the rifampicin-induced persisters are tolerant to multiple classes of antibiotics. We next tested the metabolic activity of the rifampicin-induced persisters via flow cytometry and the metabolic dye redox sensor green (RSG), which is an indicator of bacterial reductase activity and, like natural persisters (Hobby *et al.*, 1942), we found that the rifampicin-induced persister cells are metabolically inactive **(Supplementary Figure S3)**. In addition, we found there was no change in the minimum inhibitory concentration for the rifampicin-induced cells compared to the wild-type strain **(Supplementary Table 1 and Supplementary Figure S4)**, as is expected for persister cells (Brauner *et al.*, 2016). Like natural persisters (Keren *et al.*, 2004), we found that there are no mutations after the rifampicin pretreatment step and after ampicillin exposure to remove non-persister cells since re-growth of these cells led to a population that is as susceptible to ampicillin treatment as the wild-type strain **(Supplementary Table 1 and Supplementary Figures S5 and S6)**. Also, we investigated the rate of killing of the rifampicin-induced persister cells and found they have a slow rate of death (**Supplementary Figure S7A**), like natural persister cells (Harms *et al.*, 2016). Including the period of pre-treatment with rifampicin, we also confirmed that rifampicin-induced persister cells have a bi-phasic killing phenotype (**Supplementary Figure S7B**). Furthermore, natural persister cells easily convert from dormant cells to growing cells when they are contacted with a carbon source without antibiotic (Harms *et al.*, 2016), and we show below that rifampicin-induced persister cells both resuscitate like natural persisters and have the same morphology. Based on these seven independent lines of evidence, these rifampicin-induced persisters are bona fide persister cells.

### Persister cells revive immediately but are not synchronized

To avoid artifacts related to heterogeneity of cell waking, we chose to observe single cells waking on an agarose pad using light microscopy (**Figure 1**, Supplementary Movies S1 and S2) rather than monitoring growth of planktonic cells; in this manner, we could detect differences in the cell populations in their waking patterns. Furthermore, given the low number of natural persister cells, it was not possible to monitor planktonic cell growth as many hours passed before measurable changes in turbidity occurred.

For exponential cells, cell growth was seen immediately as cell division on the agarose pad (Figure 1A). After one hour, two to three cell divisions are clearly discerned, indicating cells grow well on the agarose pad. By three hours, microcolonies started to develop on the agarose pad.

**Fig. 1.**
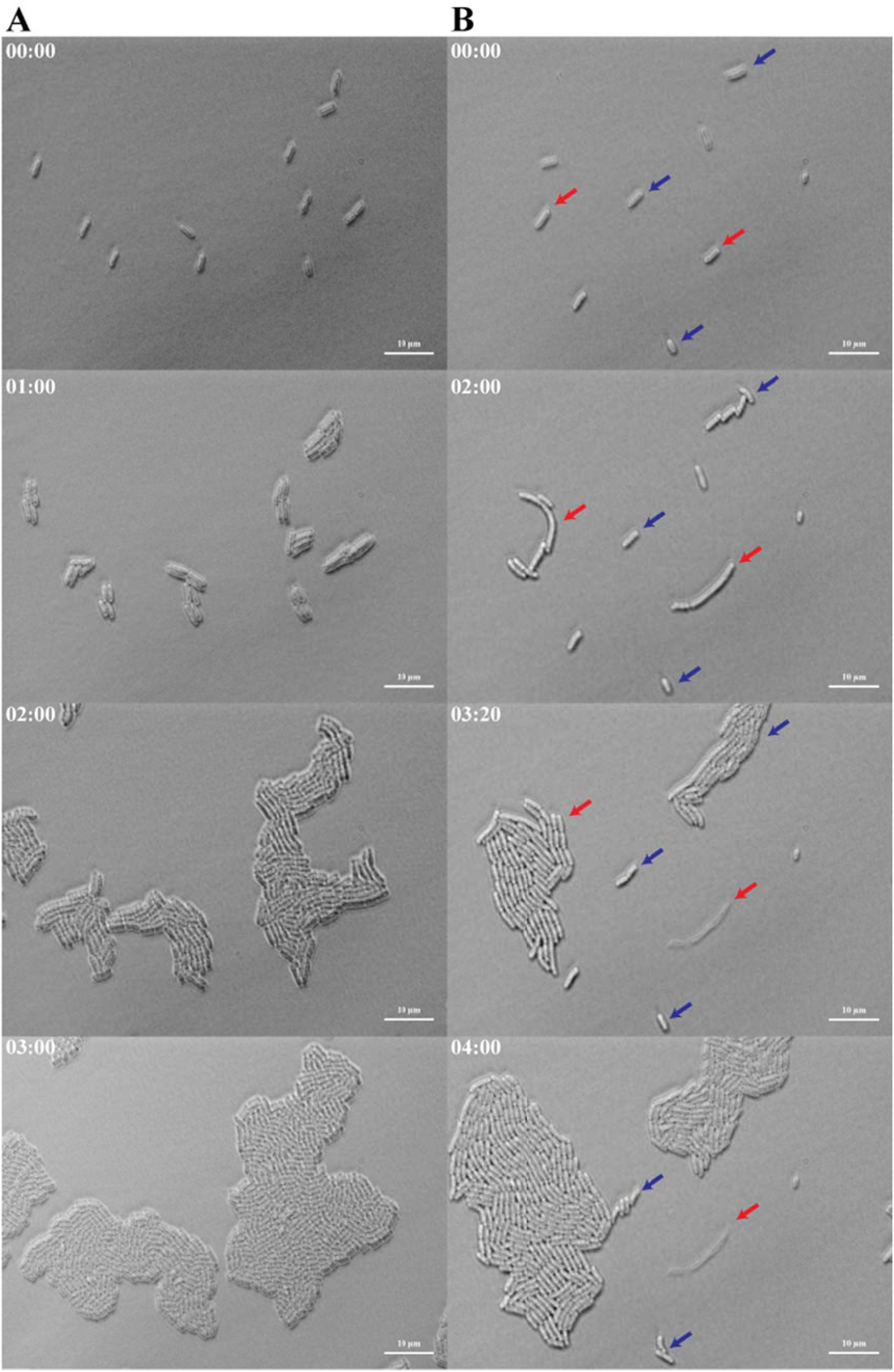
Persister waking on agarose gel pads. (**A**) Exponential-phase cells growing on agarose gel pads. All cells grow immediately by cell division. (**B**) Rifampicin-induced persisters waking on agarose gel pads (elongation indicated by red arrows and cell division indicated by blue arrows). Cell growth on agarose gel pads was visualized via light microscopy with times indicated in the upper left hand side for each panel. Scale bar indicates 10 µm.

For the rifampicin-induced persister cells (based on observation of 49 single cells), they initially showed the same morphological characteristics of exponential cells in that they have a rod-shape and are about 2 µm (Figure 1B). However, unlike exponential cells, during persister waking, five phenotypic variants were observed. Thirty-six (73%) of the cells grew and 13 (27%) failed to grow in 24 h (Table 1, Figure 2A). Of the 36 growing cells, two groups of cells began to grow immediately: two cells (4%) began to divide immediately (Figure 2B) and five cells (10%) started elongating immediately. Hence, persister cells are capable of growing immediately once the antibiotic is removed and nutrients are provided. Of these five cells that elongated immediately, three failed to divide during the six hours of observation, and two cells eventually died (Figure 2C). Of the remaining 29 cells (59%), 24 of the cells had a delay before cell division, and these persister cells showed the same phenotype as the exponential cells except for the lag time. After the delay, five of these 29 cells elongated, then divided (Figure 2D).

**Table 1.**
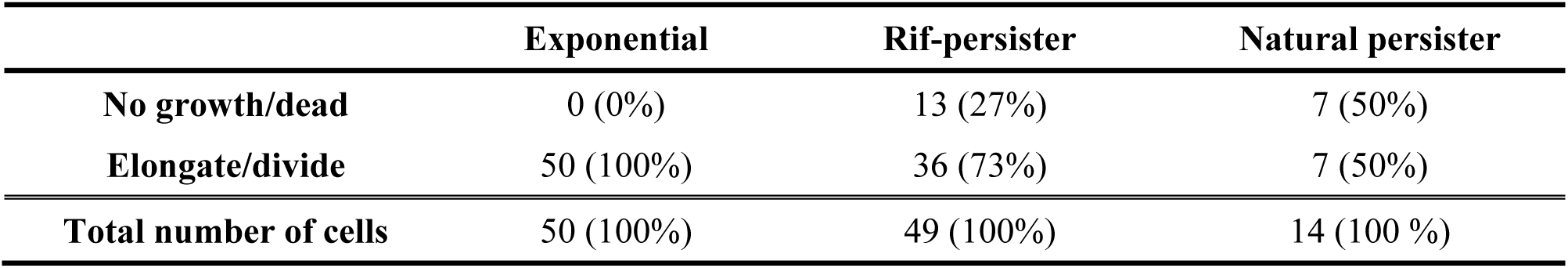
Single cell growth of exponential cells, rifampicin-induced persisters (Rif-persister), andnatural persisters on agarose pads.

**Fig. 2.**
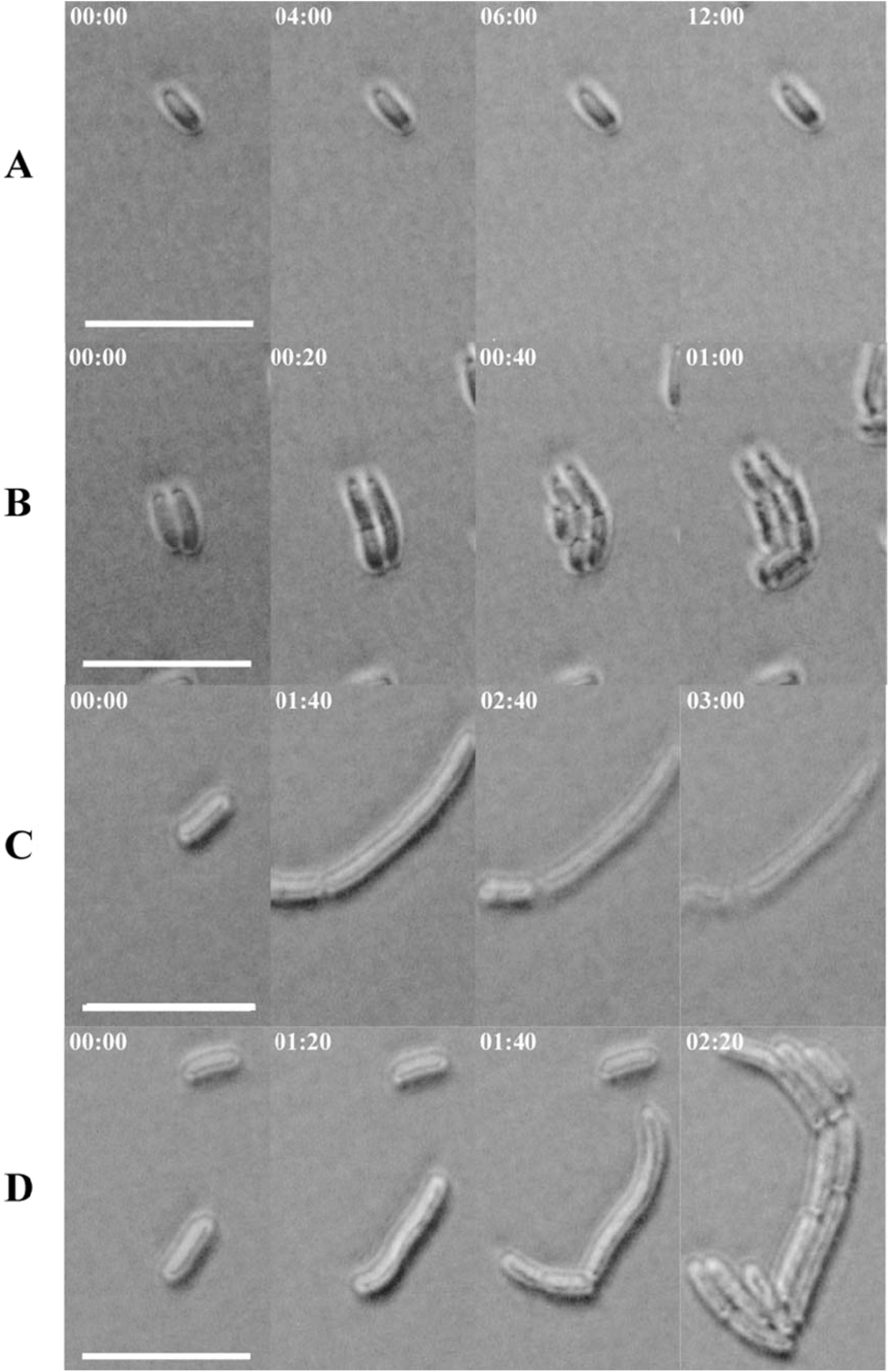
Phenotypic heterogeneity of persister cell waking. **(A**) Rifampicin-induced persister cell notchanging (for up to 24 hours). (B) Rifampicin-induced persister cell dividing without elongation. Rifampicin-induced persister cell elongating then dying (becomes transparent). (D) Rifampicin-induced persister cell elongating then dividing. Most daughter cells from this phenotype were elongated compared to exponentially-growing cells. Cell growth on agarose gel pads was visualized via light microscopy with times indicated in the upper left hand side for each panel. Scale bar indicates 10 µm.

We also examined the fate of individual natural persister cells on the agarose pads (**Supplementary Figure S8**). These cells are difficult to examine, of course, due to their low cell numbers. Critically, of the seven cells that were examined and had growth, six of them began growing immediately and one began elongating (Table 1). Also, 50% of the natural persister did not grow in 24 hours. Furthermore, two cells initiated elongation after 20 hours of contact with fresh medium that lacks antibiotics which means that they have extremely long adaptation times. Therefore, even though the population is small, these results corroborate our results with the rifampicin-induced persister cells. Hence, some persister cells resume growth immediately.

### Persister growth rates

We also investigated the growth rate of persister cells upon waking by tracking 36 single cells using time-lapsed microscopy. Only persister cells that resuscitated within 6 h were analyzed and only cell division (not cell elongation) was quantified.

For exponential cells, each cell grew immediately on the gel pad and showed the same growth rate (Figure 3A). Unlike exponential cells, the rifampicin-induced persisters showed various lag-times (including cell elongation for some cells) before cell division commenced (Figure 3B). However, each persister cell had the same growth rate as the exponential cells once cell division started (Figure 3C). This result shows clearly that once cell division occurs, persister cells return immediately to exponential growth. Similar to the rifampicin-induced persisters, after waking, the, natural persister population showed the same doubling time (21 ± 1 min) as exponential cells (Figure 3C). These observations indicate that persister cell waking is not synchronized.

**Fig. 3.**
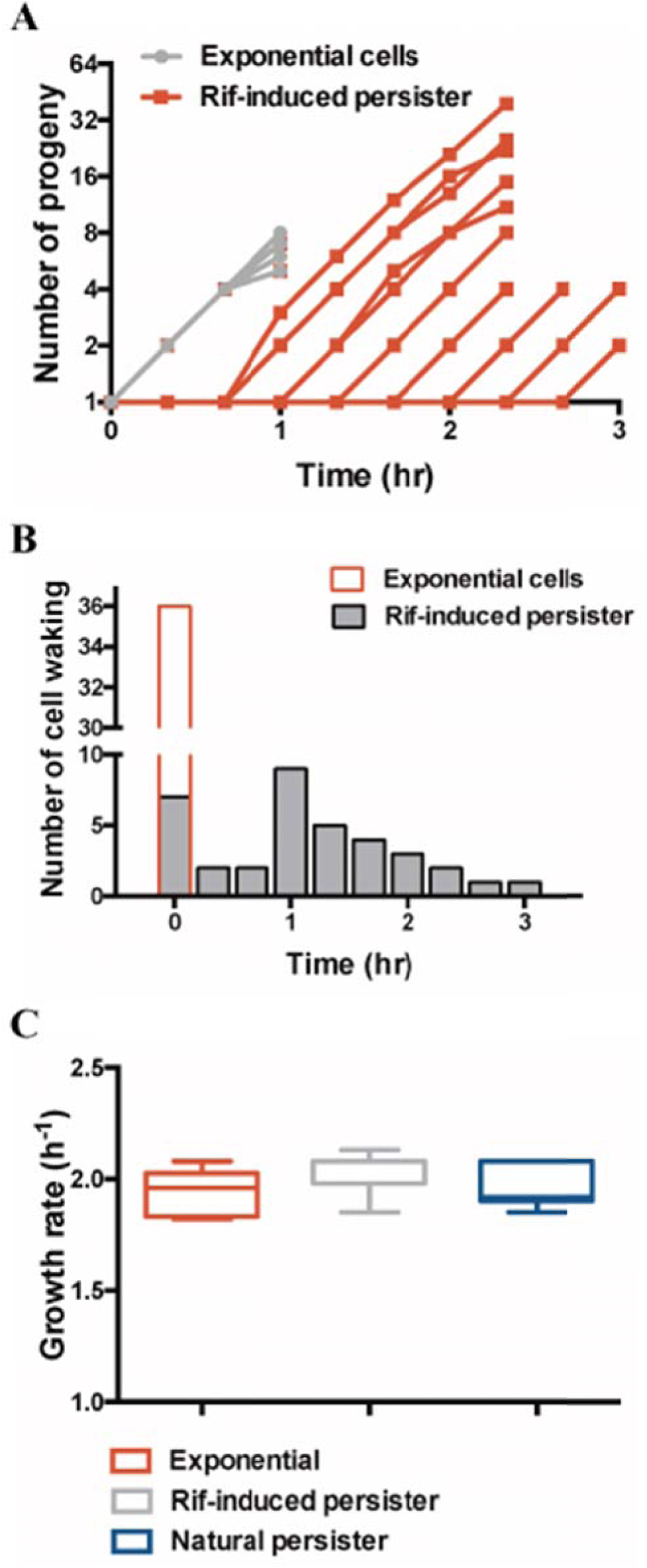
Persister waking is not synchronized. (**A**) Single cell tracking for rifampicin-induced persister cell doubling on agarose gel pads (note some cells elongate before doubling but only doubling is tracked). (**B**) Time course of the number of persister cells and exponentially-grown cells waking. (**C**) Specific growth rates of exponential cells, rifampicin-induced persister cells (“Rif-induced”), and natural persister cells.

### Active ribosome content and persister waking time

Since persister wake times were not synchronized (**Figure 3**), since cells demonstrated different morphologies upon waking (**Figure 2**), and since bacteria are 50% protein on a dry cell basis, we hypothesized that waking could be a function of active ribosome content. Under exponential growth, bacterial populations are somewhat stochastic in transcription and translation (Avery 2006); hence, ribosome content prior to antibiotic treatment should vary.

To address this hypothesis, we monitored the growth of persister cells of *E. coli* MG1655-ASV that synthesizes an unstable variant of GFP (half-life less than 1 h) as a function of ribosomal promoter activity (*rrnb*P1) (Shah *et al.*, 2006). The *rrnb*P1 promoter controls an operon of *rrsB* (16S rRNA), *gltT* (tRNA-glu), *rrlB* (23S rRNA) and *rrfB* (5S rRNA), which encode the three major rRNA building blocks of ribosomes. Since a single ribosome consists of many subunits (e.g., 35 subunits for the 50S ribosomal subunit and 23 subunits for the 30S ribosomal subunit), direct counting of active ribosomes (70S) in single cells is not easy nor convenient. Even though the measurement of ribosomal promoter activity is not a direct observation of ribosome number, using this method as a proxy for the number of ribosomes in the cell based on measurement of rRNA concentration has been used in many papers (Burger et al., 2010, Lu *et al.*, 2009, Piques *et al.*, 2009). To confirm that the GFP reporter in this strain indicates ribosome levels, we assayed the fluorescence of single cells in exponential (turbidity 0.8 at 600 nm) vs. those in stationary growth (turbidity 4.5 at 600 nm); ribosomes are produced at higher concentrations at higher growth rates (Bremer and Dennis 2008). As shown (**Supplementary Figure S9A**), single-cell fluorescence is clearly more pronounced (approximately more than 10-fold) in exponential cells relative to stationary-phase cells. To further support the use of the GFP reporter as a proxy for active ribosome levels, we correlated active ribosome levels in exponential, rifampicin-induced persister, and stationary-phase cultures with GFP intensity. Direct observation of active ribosome levels clearly shows that the GFP intensity is correlated with active ribosome contents in the cells (**Supplementary Figure S9B**). Therefore, the GFP signal is a robust indicator of active (70S) ribosome levels in the single cells.

Critically, we found the subpopulation of rifampicin-induced persister cells with higher GFP intensity (after ampicillin treatment) started to grow immediately on the agarose pad once nutrients were provided (**Figure 4**, Supplementary Movie S3, **red arrows**); note these cells were shown to be metabolically-inactive after rifampicin treatment **(Supplementary Figure S3**), so they wake immediately in the presence of nutrients. In the presence of antibiotics, persister cells cannot produce more protein since they are dormant; hence, the GFP intensity on the agarose pad with fresh nutrients indicates the ribosome content prior to antibiotic treatment and any possible decay of ribosomes during antibiotic treatment. To confirm this, we observed the GFP intensity of rifampicin-induced persister cells on agarose pads containing ampicillin (100 µg/mL); i.e., we observed the cells during the ampicillin treatment. At the start of the ampicillin treatment, all the persister cells have relatively high GFP signals since they were generated from exponential cells; but, some have approximately 2-fold higher fluorescence. Over the three hours of exposure to ampicillin (**Supplementary Figure S10A, red arrows**), all the cells lost GFP at the same rate (**Supplementary Figure S10B**) but since some cells started at higher GFP levels, a small percentage of cells retained significant fluorescence intensity whereas most persister cells had their GFP intensity diminish to low levels (**Supplementary Figures S10A**, blue arrows). To support this result, we confirmed that the rifampicin treatment inhibits the synthesis of new GFP by comparing the GFP signal of cells without rifampicin to those with rifampicin (**Supplementary Figures S10C**); without rifampicin, cells maintained their GFP signal while cells that were treated with rifampicin had their GFP signal decay since the rifampicin prevents protein synthesis. Based on these results, we conclude that the GFP signals originate from production of ribosomes prior to antibiotic treatment and that there may be degradation of ribosomes during the antibiotic treatment stage. Critically, for the cells with low GFP intensity at 0 min on the agarose pad, some of them increased their GFP intensity and started to grow after 120 min in the fresh medium of the agarose pad (**Figure 4**, Supplementary Movie S3, **blue arrow**). Therefore, for these persister cells formed from mid-exponential-phase cells at turbidity at 600 nm of 0.8, the cell population having high ribosome content prior to antibiotic treatment resuscitated immediately then divided (or died) while the cell population having reduced ribosome content divided (after a delay) without cell elongation (Figure 4A).

**Fig. 4.**
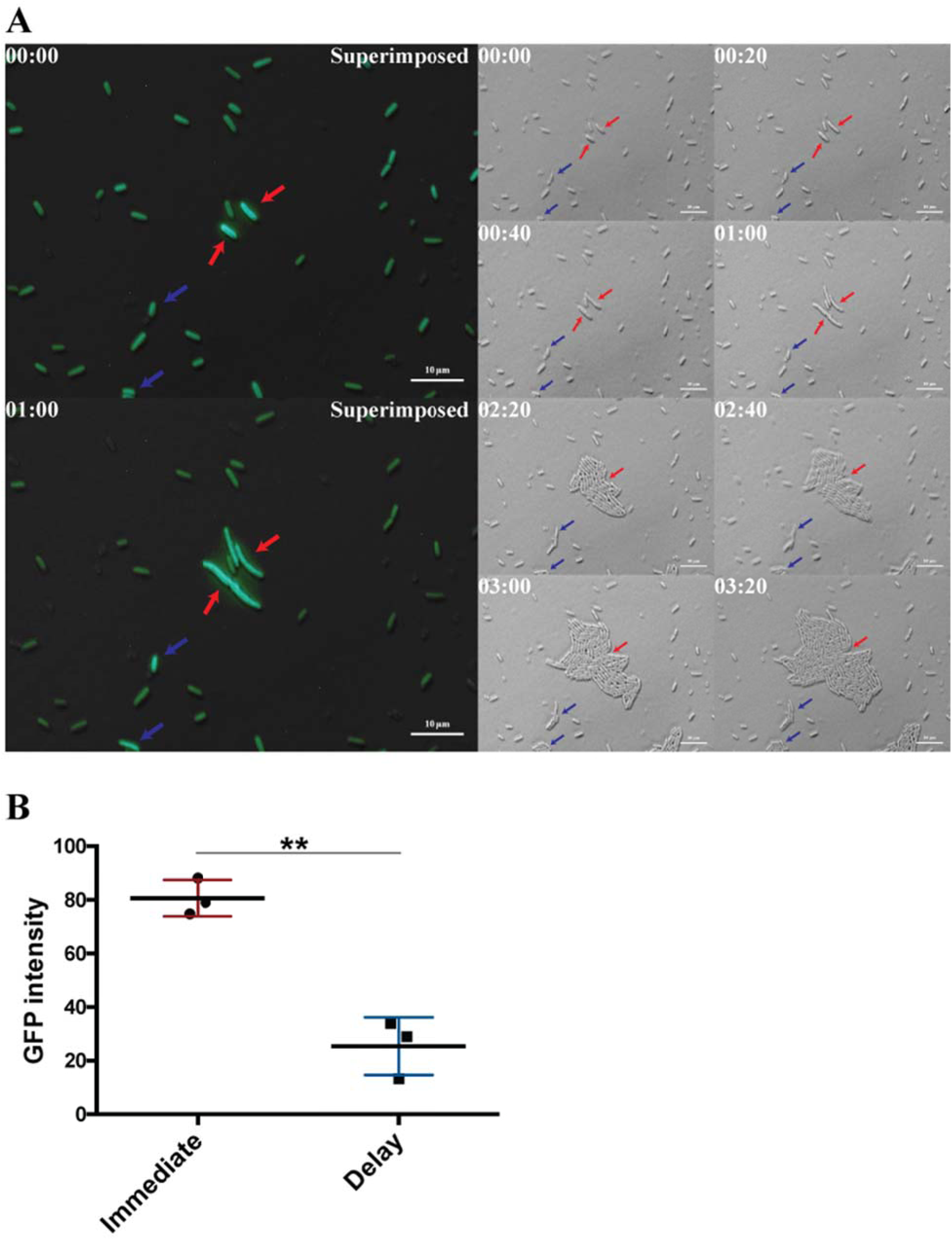
Heterogeneity of persister waking and ribosome content. (**A**) Persister cells(*rrnb*P1::GFP[ASV]) formed by rifampicin-treating exponential-phase cells with high numbers of ribosomes prior to antibiotic treatment (red arrows, left panel in first two rows) initiate cell growth (elongation) immediately when contacted with the carbon source without antibiotics. Rifampicin-induced persister cells with fewer ribosomes increase ribosome content upon contacting the carbon source (blue arrows) and have longer waking times and cell division without cell elongation. Cell growth on agarose gel pads was visualized via fluorescence microscopy (left panels) and with light microscopy (middle and right panels); times indicated in the upper left hand side for each panel. Scale bar indicates 10 µm. (**B**) Comparison GFP intensity (which is indicates ribosome content) between persister that wakes immediately and persister thatwakes after a delay. Statistical significance was determined by Student’s *t* test.**, *P* < 0.01.

We quantified the level of ribosomes based on GFP fluorescence and found the persister cells that resuscitated immediately had about 4 times more ribosomes than the persister that resuscitated after a delay (Figure 4B). Unfortunately, we could not find any differences in initial GFP intensity between persister cells that resuscitated after a delay (1 to 6 h) and persister cells that never resuscitated in 6 h; however, the persister cells that resuscitated after a delay increased their ribosome contents (GFP intensity) unlike those cells that did not resuscitate (Figure 4A).

To corroborate these findings by producing a large population persister cells with high ribosome content (so that we could discern differences in this subpopulation), rifampicin-induced persister cells showing high GFP intensity were sorted by flow cytometry based on GFP intensity. To exclude dead cells, propidium iodide (PI)-stained dead cells were removed, and the persister cells with the highest 1.25% of GFP signal were sorted directly onto LB agarose gel pads to minimize both cell loss and the time that persister cells were in PBS buffer (**Supplementary Figure S11A**). As expected, most of persister cells with high GFP intensity resuscitated immediately (**Supplementary Figure S11B**). However, not all persister cells with high GFP intensity resuscitated (**Supplementary Figure S11B, orange arrows**). Also, due to errors in sorting, a few cells were obtained with low GFP intensity; these cells with low ribosome content did not wake immediately, as expected (**Supplementary Figure S11B, red arrow)**. Corroborating these results, when rifampicin-induced persister cells with low GFP intensity were sorted directly onto LB agarose gel pads, in contrast to cells with high GFP content, no cells with low ribosome levels resuscitated immediately, and those with low ribosome (GFP signal) that did resuscitate, did so after a 120 min delay.

To further support the finding that persister cells wake based on ribosome content, we induced persister cell formation via rifampicin treatment in both the early exponential phase (turbidity at 600 nm ~ 0.3) and stationary phase (turbidity at 600 nm ~ 5) and observed the waking phenotype and ribosome content. As expected, rifampicin-induced persisters of the early exponentially-growing cells (**Supplementary Figure S12**) showed higher ribosome content than persister cells formed from stationary cells (**Supplementary Figure S13**). Also, most of the persister cells formed from the early exponential cells resuscitated immediately while the persister cells formed from stationary cells divided after a 1.3 hr delay. Also, as seen previously with the persister cells formed using mid-exponential cells that had low GFP intensity (**Figure 4**), the persister cells formed from stationary-phase cells increased their GFP intensity upon waking then divided (**Supplementary Figure S13**). Together, these results clearly show the ribosome content dictates cell division; i.e., the ribosome content must increase in persister cells to some threshold level prior to cell division which dictates whether cells elongate immediately or pause and then divide; however, high ribosome content itself does not guarantee waking.

### Persister elongation is not caused by damage of penicillin binding proteins (PBPs)

β-Lactams inhibit the PBPs that catalyze the cross-linking of peptidoglycan (Nanninga 1998) causes cell elongation (Spratt 1975); hence, we hypothesized that some persister cells may contain more PBPs which leads to cell elongation at the initiation of waking. To test this hypothesis, Bocillin-FL (which is not active as an antibiotic, **Supplementary Table S1**) was employed along with ampicillin for rifampicin-induced persisters; Bocillin-FL is penicillin with the fluorescent tag BODIFY that likely diffuses freely into the periplasmic space via OmpF (Kojima and Nikaido 2013) and detects PBPs by binding (Zhao *et al.*, 1999). Bocillin-FL does not change its fluorescence appreciably upon binding (Shapiro *et al.*, 2013). We found that both persister cells that wake by cell elongation (**Figure 5**, **red arrow**) and persister cells that wake by cell division (**Figure 5**, **blue arrow**) have no fluorescence signal with Bocillin-FL, which indicates that there is no difference in corruption by ampicillin in these persister cells since the number of PBPs is low. However, these results may be complicated by a possible competition between Bocillin-FL and ampicillin in the rifampicin-induced persister culture. Furthermore, some un-lysed cells had a strong Bocillin-FL signal and failed to resuscitate in 6 h (**Figure 5**, **pink arrow**); these might be dead cells that are more permeable. Therefore, for the persister cells that resuscitate, the elongation phenotype seen appears to not be due to ampicillin corruption of PBPs since there is no detectable Bocillin-FL in these persister cells; however, high PBP concentrations in the persister cells leads to no resuscitation. As a positive control, exponential cells treated by Bocillin-FL were uniformly stained by Bocillin-FL (**Figure 5**, **upper left panel**). As a negative control, stationary-phase persister cells did not fluoresce like persisters in exponential culture (**Figure 5**, **upper right panel**).

**Fig. 5.**
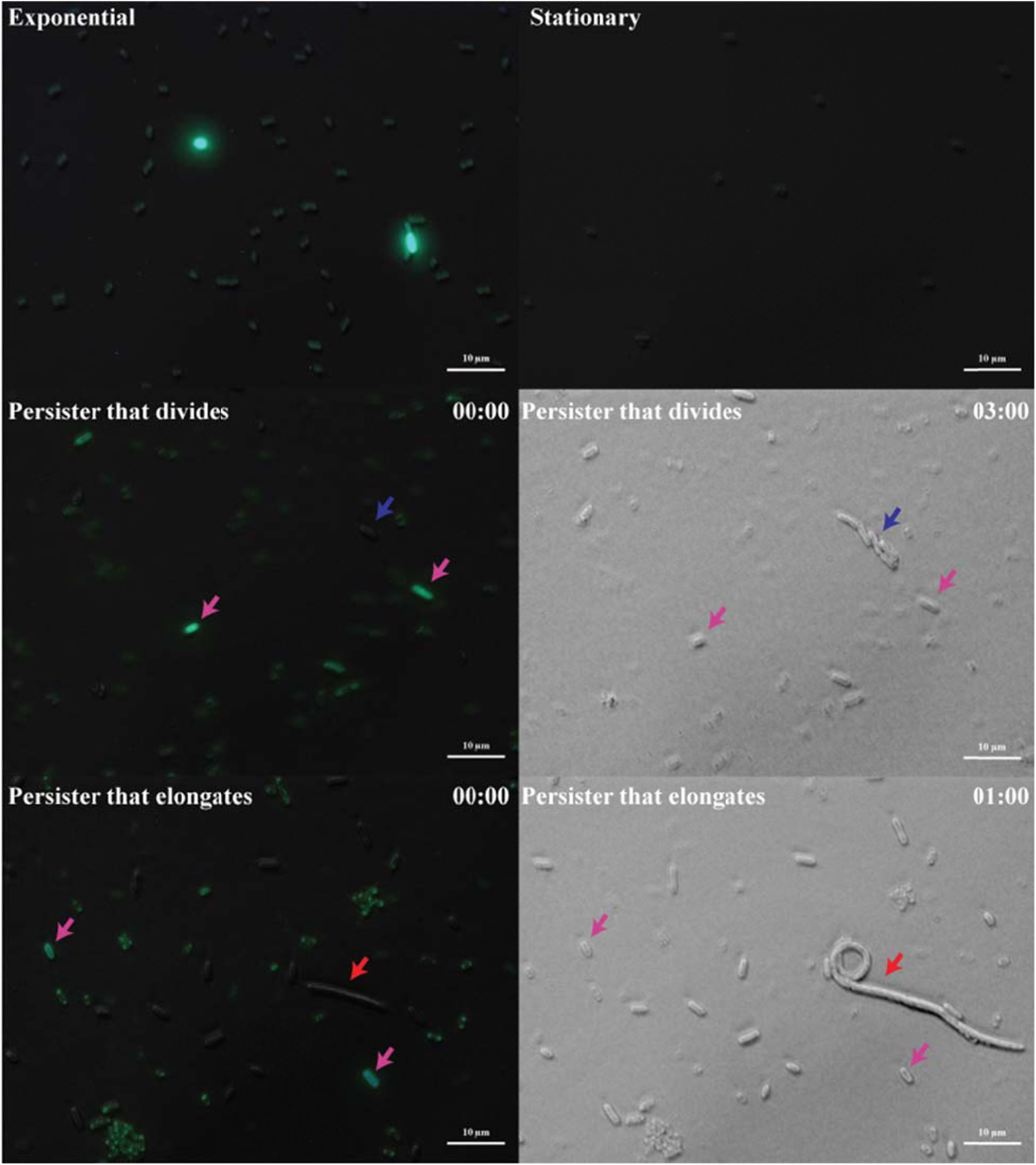
Residual ampicillin in exponential, stationary-phase, and persister cells. Superimposed images combining bright and BODIFY channels (green). Exponential cells with uniform Bocillin-FL signal (upper left panel), stationary-phase cells with no Bocillin-FL signal (upper right panel), and rifampicin-induced persister cells (middle left and middle right panels) that (i) divide and lack Bocillin-FL signal (blue arrow), (ii) that have Bocillin-FL signal but do not wake (pink arrows), and (iii) that elongate and lack Bocillin-FL signal (red arrow).

## DISCUSSION

By the seven lines of evidence (muti-drug tolerance, easy conversion from persister state to non-persister state, dormancy, no change in MIC compared to exponential cells, no resistance phenotype, similar morphology to natural persisters, and similar resuscitation as natural persisters) (**Supplementary Figures S2 to S8),** our results confirm that rifampicin-induced persister cells (Kwan *et al.*, 2013) are equivalent to those that arise naturally without pretreatment. Furthermore, by making the persister cell phenotype dominant through rifampicin pretreatment, we discovered, through microscopy of individual cells, that persister cell resuscitation is not synchronized in that there are five different cell types of persister cell resuscitation (**Figure 6**). Critically, some of these antibiotic-tolerant persister cells resuscitate immediately, revealing that persister cells respond immediately to the removal of antibiotics and to the addition of fresh medium. This indicates that any procedure such as a wash with fresh medium (rather than non-nutritive buffer) immediately wakes some persister cells and invalidates the subsequent results as related to persistence (Orman and Brynildsen 2013, Pu *et al.*, 2016). Moreover, this non-synchronized resuscitation of persister cells suggests the phenotypic heterogeneity prior to persister cell formation manifests itself upon persister waking. Also, in both the rifampicin-induced and natural persister populations, non-resuscitated cells were found among the unlysed cells when the antibiotic stress was removed. These cells may be dead since not all dead cells lyse, but we counted these cells as persisters since they are not lysed in presence of ampicillin; hence, our estimate of the number of persister cells may be somewhat elevated.

**Fig. 6.**
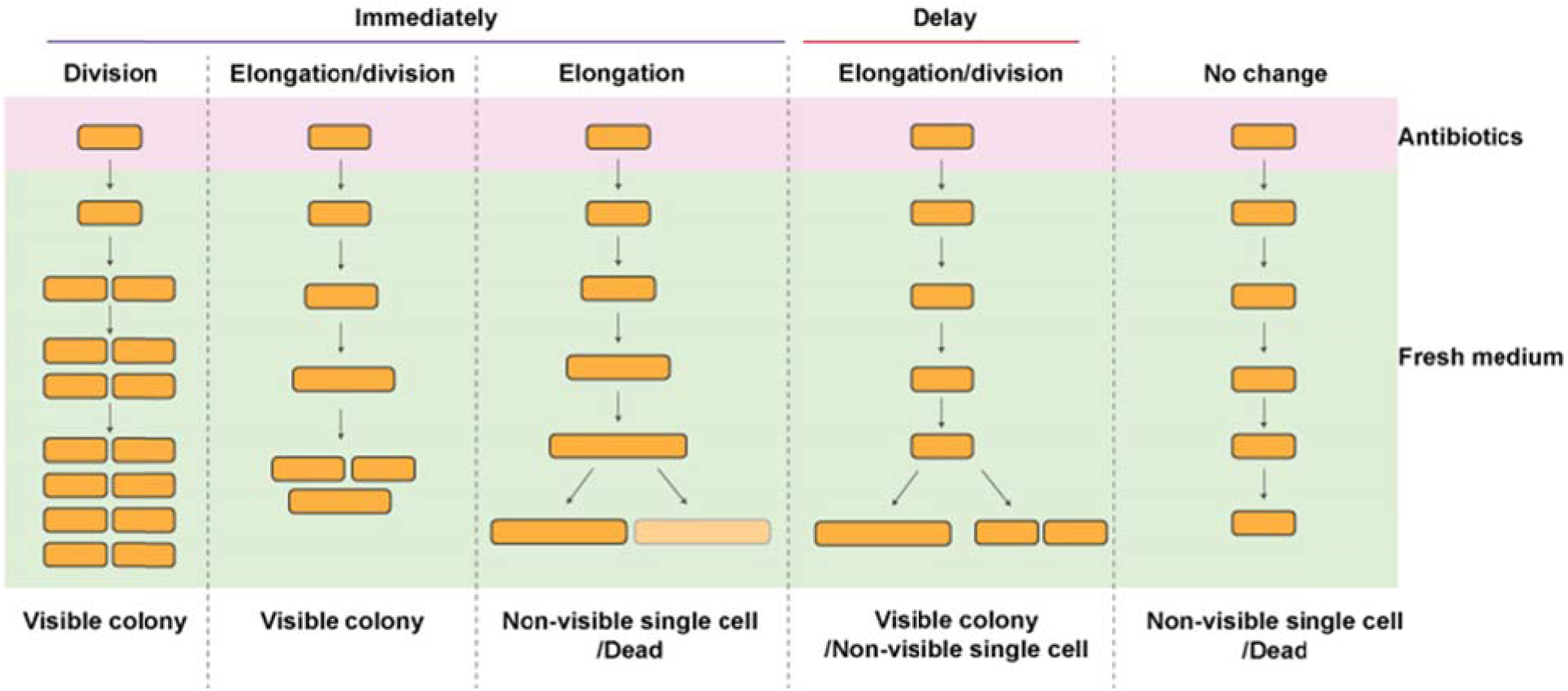
Model for morphological dynamics upon persister waking. The unlysed (persister) cells after ampicillin treatment had five phenotypes upon contacting fresh media. The cell types that divide showed various waking times, and the other cell types showed only elongation. Some un-lysed cells showed no resuscitation even after the antibiotic stress was removed.

The elongation of bacteria during antibiotic exposure has been documented by many researchers (Chung *et al.*, 2009, Martinez et al., 1982, Yao *et al.*, 2012); however, the elongation phenotype seen with the persister cells in our work occurred after the antibiotic was removed (Figure 2CD). Interestingly, the daughter cells from this persister population were elongated for several steps of cell division unlike exponential cells. For example, the daughter cells from the exponentially-grown culture were 2 µm in average length (same as their maternal cell) but the daughter cells from persister elongated cells were about 5 µm in average length (Figure 2D). Furthermore, the elongation followed by death phenotype seen with some persister cells (Figure 2C) indicates that classic methods to determine persister cell activity do not fully reflect the short-term viability of the whole persister population as these cells would fail to form a colony. Moreover, the elongation of the rifampicin-induced persister cells on agarose pads during resuscitation is similar to the cell elongation seen previously in a persister cell with high guanosine tetraphosphate levels (Maisonneuve *et al.*, 2013) and persister cells formed by production of toxin MazF of the MazF/MazE toxin/antitoxin system (Cho *et al.*, 2017); this latter phenotype was seen with ciprofloxacin. Combined with our observations, cell elongation during persister waking is a common phenotype that is not restricted to the use of ampicillin to form persister cells via corruption of cell wall synthesis. Furthermore, our observations of PBP levels in persister cells via Bocillin-FL labelling indicates that persister cell elongation is not due to more corruption of PBPs by ampicillin since both persister cells that elongate and divide have no differences in Bocillin-FL signal (**Figure 5**); hence, these persister cells have low concentrations of PBPs. This is reasonable since compared to exponential cells, we found stationary-phase cells did not show Bocillin-FL fluorescence, and stationary-phase cells are known to have reduced levels of PBPs (Stevens *et al.*, 1993). Critically, persister cells with high Bocillin-FL levels showed no resuscitation in 6 h (**Figure 5**, **pink arrow**), so the corrupted PBPs prevent resuscitation.

Once the persister cells (that do not elongate first) begin to divide, they grow with the same doubling time as exponential cells (21 min, Figure 3AC) which means that once division begins for these cells, the persister cell is fully recovered. However, the time required for the persister cells to begin to divide varies (Figure 3A) and reflects heterogeneity of the cells prior to persister cell formation; i.e., it indicates there is a different level of cell metabolism upon waking from the dormant state. This difference in metabolism was found here to be dictated not by antibiotic content but by the ribosome content; i.e., cells with higher numbers of ribosomes resuscitated immediately (Figure 4A and **Supplementary Figure S11**). Also, our results indicate that heterogeneity of the cell population before persister cell formation is preserved in the persister state. The persister cells that wake immediately use ribosomes that existed prior to persister cell formation while the persister cells with delayed waking needed to produce ribosomes to revive cellular metabolism (Figure 4A and **Supplementary Figure S11**). However, ribosome content of the persister cell alone does not dictate whether persister cells wake or not (Figure 4A, **Supplementary Figures S11, orange arrow**) since not all persister that have low ribosome content or high ribosome content resuscitated in 6 hour with a fresh carbon source and the antibiotic removed. Critically, since we have shown cessation of translation makes cells become persisters (Kwan *et al.*, 2013), it makes sense that the number of active ribosomes should control the way cells wake. Nevertheless, how persister cells sense fresh medium that lacks antibiotics requires further study.

## ACKNOWLEDGMENTS

This work was supported by the Army Research Office (W911NF-14-1-0279) and funds derived from the Biotechnology Endowed Professorship at the Pennsylvania State University. We thank Professor Kim Lewis for *E. coli* K-12 MG1655-ASV.

## SUPPLEMENTARY VIDEO LEGENDS

**Supplementary Movie S1. Exponential-phase cell growth on an agarose gel pad (3 h).** Exponential-phase cells growing on an agarose gel pad at 37 ºC demonstrating all cells grow immediately by cell division. Images were taken every 20 min, and the scale bar indicates 10 µm.

**Supplementary Movie S2. Persister resuscitation on an agarose gel pad (4.3 h).** Resuscitation of rifampicin-induced persister cells on an agarose gel pad at 37 ºC demonstrating morphological dynamics that include (i) waking by elongation followed by cell division, (ii) waking by elongation followed by cell death, (iii) waking by cell division without cell elongation and (iv) no waking. The red arrows indicate the persister cell that grow immediately by elongation. The blue arrows indicate the persister cell that increase ribosome contents then grow by cell division without elongation. Images were taken every 20 min, and the scale bar indicates 10 µm.

**Supplementary Movie S3. Persister resuscitation of GFP-labeled *E. coli* on an agarose gel pad (5 h).** Resuscitation of rifampicin-induced persister cells on an agarose gel pad at 37 ºC demonstrating that cells with high ribosome content (**Figure 4**, **red arrow**) grow immediately and demonstrating cells with low ribosome content (**Figure 4**, **blue arrow**) increase ribosome content first then grow by cell division without elongation. The red arrows indicate the persister cell that grow immediately by elongation. The blue arrows indicate the persister cell that increase ribosome contents then grow by cell division without elongation. Images were taken every 20 min, and the scale bar indicates 10 µm.

